# Microbial diversity in raw milk and Sayram Ketteki from southern of Xinjiang, China

**DOI:** 10.1101/2021.03.15.435442

**Authors:** DongLa Gao, Weihua Wang, ZhanJiang Han, Qian Xi, RuiCheng Guo, PengCheng Kuang, DongLiang Li

**Author notes:** Corresponding author (Weihua Wang).

## Abstract

Raw milk and fermented milk are rich in microbial resources, which are essential for the formation of texture, flavor and taste. In order to gain a deeper knowledge of the bacterial and fungal community diversity in local raw milk and home-made yogurts from Sayram town, Baicheng county, Akesu area, southern of Xinjiang, China,30 raw milk and 30 home-made yogurt samples were collected and experiment of high-throughput sequencing was implemented.The results of experiments revealed the species of fungi in raw milk was the most, and the species of bacteria in fermented milk was the least.Based on principal component analysis (PCA), it was found that the bacterial and fungal community structure differed in samples from two types of dairy products.And the presence of 15 bacterial and 12 fungal phyla, comprising 218 bacterial and 495 fungal genera respectively, among all samples. Firmicutes and Ascomycota,Lactobacillus and Candida were the predominant phyla and genera of bacteria and fungi, respectively. The results indicated that the microbial community of raw milk differs from home-made yogurts due to sampling location and manufacturing process. The study suggested that high-throughput sequencing could provide a better understanding of microbiological diversity as well as lay a theoretical foundation for selecting beneficial microbial resources from this natural yogurt.

## 1. Introduction

Raw milk (RM) and dairy products derived therefrom are rich in nutrients and microorganisms, and their microbial composition have been discovered in previous studies, through traditional culture-dependent and culture-independent methods [1,2].Microbial differences in RM are related to many factors, such as animal sources and geographic locations. In the flora of 14 pastures, the leading bacteria genera were *Enterococcus*, *Bacillus*, *Acinetobacter* and *Lactococcus* but only *Staphylococcus* aureus and *Shigella* were detected in a few cattle farms [3]. In donkey milk, it was found that the dominant bacterial phyla were Proteobacteria and Firmicutes with dominant genera being *Pseudomonas*, *Ralstonia* and *Acinetobacter*[4]. In goat milk, it was found that the dominant genera were *proteobacteria* and *Enterobacter* among 354 genera[5].

Generally, the most dominant bacterial genera in dairy products are *lactobacillus*, *lactococcus*, *streptococcus* and *leuconostoc* of Firmicutes, and they affect the flavor and maturity of fermentation dairy products [6, 7]. For example, lactic acid bacteria play an important role in the fermentation process, in which they heterogeneously ferment to generate ethanol, carbon dioxide, acetaldehyde, acetone, acetoin, diacetyl and other volatile aromatic compounds, promoting the formation of good food flavors[8].Yeast fermented dairy products, such as kefir, koumiss and mozzarella, are typical fermented milk with alcohols, which are important contributors to fragrance and wine aroma [9]. It was revealed that *Geotrichum candidum* was a main contributor to the maturation of camemberti cheese and had the ability to metabolize amino bitter peptides and enhance the sulfur taste[10].

Although traditional cultivation methods and molecular technologies made us have a preliminary understanding of microbial composition in dairy products [11,12], the information obtained is not always accurate and the details of these products’ microbial diversity remains elusive. The main reason is that a large number of microbial species could not be cultured successfully [13].

In the recent years, with the mutual penetration of genetics, bioinformatics and other disciplines, more and more culture-free technologies have been used in the studies of microbial diversity[14]. Metagenomics[15] is a fast and efficient method which uses high-throughput sequencing to simultaneously sequence millions of DNA molecules, and constructs a metagenome library by detecting the genetic material sequences of variable regions in microorganisms, exploring the diversity of microbial species and community structure. This technology avoids non-cultivable and low-abundance microbial unpredictability, and objectively restores the community structure and abundance ratio, maximizing the development of microbial resources. This approach has been widely used in medical and environmental science [16, 17], and gradually used in the food science[18].

Through metagenomics, the diversity of microorganisms, dominant flora and metabolic potential in Mexican Cotija cheese was discovered[19]. It indicated that the *Lactobacillus, Leuconostoc* and *Weissella* were the main dominant flora, the microbial metabolic activities related to the formation of multiple flavors mainly were involved the metabolism of branched chain amino acids and free fatty acids, and also discovered some genes related to bacteriocin production and immunity. *Shewanella, Acinetobacter*, *Pelomonas*, *Dysgonomonas*, *Weissella* and *Pseudomonas* were found in Tibetan kefir grains for the first time[20]. In Turkish Kefir grains, it was found that communities in the two samples showed high consistency. *Lactobacillus* including *L. kefiranofaciens*, *L. buchneri* and *L. helveticus* was the most abundant genus[21]. The diversity of fungi in French Tomme d’Orchies cheese was interesting. The results showed that it contained *Yarrowia lipolytica* and *Galactomyces geotrichum*, and some rare microorganisms had been discovered, such as *Clavis-poralusitaniae*, *Kazachstania unispora* and *Cladosporium cladosporioides*[22].

The microbial diversity of fermented dairy products, such as koumiss[23], jueke [24], tarag [25]and airag [26], had been investigated widely, but there were few reports on fermented dairy products in Xinjiang province of China [27], especially in south Xinjiang. Sayram Ketteki (SK), a traditional and popular handcraft fermented food, was produced by local Uighur peasant woman in Xinjiang province of China. To generate SK, the milk from cows is used as the raw material, and a little yogurt reserved in the previous batch acts as starter of fermentation. Because of continuous making yogurt, the fermentation starters from some families even have decades of history. After fermenting in porcelain bowels at 10-25℃ about 6-8 hours, it obtains the products with the color of milky white, elastic, sweet and sour-tasting with a slightly alcoholic. SK was typical alcoholic fermented milk. Several studies have investigated the lactic acid bacteria and yeasts in SK using culture-based method and molecular biological identification. It was reported that the dominant strains of SK were *L.Bulgaricus*, *Streptococcus thermophilus* and *Saccharomyces cerevisiae*[28]. But some others studies demonstrated that *L. helveticus*, *Streptococcus thermophilus* and *Kluyveromycus marxianus* were the predominant [29]. Although there are some related studies[30], the mechanism of its flavor formation is still unclear. Undoubtedly, the formation of flavor has a lot to do with microorganisms, therefore, unraveling microbial community diversity of SK and its raw materials is very necessary.

In the current study, to explore the microbial species diversity and the microbial composition in RM and SK, 30 RM and 30 home-made yogurt samples were collected, metagenomic sequencing were carried out. Through bioinformatics analyses, it was found that microorganisms in both types of dairy products were various. Here we report the results.

## 2. Materials and Methods

### 2.1 Sample collection and DNA extraction

Sixty samples were collected aseptically from different national minority families of Sayram town Baicheng county which located in the southern of Xinjiang (approx 41°48’ N, 81°87’ E) of China, where the average temperature was 27°C in late July 2019. RM samples were named with the letter “X” (X1, X2…X34), while SK samples were named with the letter “S” (S1, S2…S34), respectively. Each RM sample and each SK sample was corresponding, because they came from the same family. These collected samples were stored at −20℃ for further use.

Every sample was centrifuged at 10000 rpm for 10 min. The supernatant was discarded and the precipitate was left. Total microbial DNAs were extracted from 500 mg sediment for each sample using the DNeasy Power Soil Kit (QIAGEN, Netherlands), according to instructions of the manufacturer, and kept at −20 °C prior to future analysis. The quantity and quality of extracted DNAs were measured by NanoDrop ND-1000 spectrophotometer and 0.8% agarose gel electrophoresis.

### 2.2 PCR amplification of 16SrDNA and ITS sequences

The bacterial V3-V4 of 16SrRNA gene was amplified, the used primers were as follows: 338F: 5’-ACT CCT ACG GGA GGC AGCA-3’ (forward) and 806R: 5’-GGA CTA CHV GGG TWT CTA AT-3’ (reverse). For fungal community analysis, the internal transcribed spacer (ITS) gene was amplified using the primers: ITS5F: 5’-GGA AGT AAA AGT CGT AAC AAG G-3’ (forward) and ITS1R: 5’-GCT GCG TTC TTC ATC GAT GC-3’ (reverse). In the PCR experiments, the thermal cycling consisted of an initial denaturation at 98°C for 5 min, 25 cycles of denaturation at 98 °C for 30 s, annealing at 52 °C for 30 s, and extension at 72 °C for 1 min, followed by a final extension at 72 °C for 5 min.

PCR products were measured with 2% agarose gel electrophoresis, and the target fragments were collected by using gel recovery kit (AXYGEN, Dalian, China). Referring to the preliminary quantitative results of electrophoresis, the obtained products were quantified with fluorescence and according to the quantitative results and sequencing volume requirements of each sample. Subsequently, each sample was mixed in corresponding proportion and paired-end 2×300 bp sequencing was carried out using the Illumina MiSeq platform with MiSeq Reagent Kit v3 by Shanghai Personal Biotechnology Co., Ltd (Shanghai, China).

### 2.3 Sequence quality control

In order to fulfill the requirements of subsequent analyses, the all readings were screened and filtered using Quantitative Insights into Microbial Ecology (QIIME) (V1.8.0) software [31]. Briefly, the raw reads with exact matches to the barcodes were assigned to respective samples and identified as valid sequences. The low-quality sequences that were less than 150 bp, average Phred scores of less than 20, ambiguous, and contained mononucleotide repeats more than 8 bp were discarded. Paired-end reads were assembled using FLASH[32]. After chimera detection, the remaining high quality sequences were clustered into operational taxonomic units (OTUs) at 97% sequence identity with UCLUST[33]. The highest sequence was taken as the representative sequence of the OTU. OTU taxonomic classification was conducted by BLAST searching and aligning the Prokaryotic sequences with the Greengenes database[34], and Eukaryotic sequences with the Unite database[35].According to the number of sequences included in each sample of each OTU, a matrix file of the abundance of OTU in each sample was constructed.

### 2.4 Bioinformatics and statistical analysis

Sequence data analyses were mainly conducted using QIIME and R packages (V3.2.0). OTU-level alpha diversity indices, such as Chao1 and ACE richness index, Shannon and Simpson diversity index, were calculated using the OTU table in QIIME. OTU-level ranked abundance curves were generated to compare the richness and evenness of OTUs among samples. Beta diversity analysis was performed to investigate the structural variation of microbial communities across samples visualized via Principal component analysis (PCA). The taxonomy compositions and abundances were visualized using GraPhlAn [36], and correlations with |RHO| > 0.6 and P < 0.01 were visualized as co-occurrence network using Cytoscape. Microbial functions were predicted by PICRUSt (Phylogenetic investigation of communities by reconstruction of unobserved states) [37].

## 3. Results

### 3.1 Bacterial and fungal sequence richness and diversity

After filtering, we obtained 4,092,576 high-quality reads for bacteria, ranging from 60,078 to 75,419 with an average of 68,210 ± 4,393 sequence reads per sample. Meanwhile, 4,056,820 of ITS sequence reads were generated for the fungal community, ranging from 60,253 to 75,925 with an average of 67,614 ± 4,665 sequence reads per sample. The number of unique and classifiable representative OTUs sequences for bacteria and fungi were 2,221 and 4,062, respectively.

The 16SrRNA of bacteria from SK had the largest number of sequences, followed by the number of ITS sequences of fungi from RM and 16S rRNA of bacteria from RM and the number of ITS sequences of fungi from SK was the smallest (Table 1). From the Chao1 index, the highest abundance of bacteria was in RM samples, followed by bacteria of SK, fungi of RM, and the lowest abundance was fungi of SK. From the Shannon index, the highest diversity was fungi of RM, followed by bacteria of RM and fungi of SK, and the bacteria of SK had the lowest diversity. The results of the numbers of OTUS were consistent with the results of diversity, which indicated that the species of fungi in RM was the most, and the species of bacteria in SK was the least.

**Table 1:** Alpha diversity values in samples.

| sample |  | Trimmed tags |  | Chao1 index |  | Shannon index |  | Number of OTUs |  |
| --- | --- | --- | --- | --- | --- | --- | --- | --- | --- |
|  |  | bacteria | fungi | bacteria | fungi | bacteria | fungi | bacteria | fungi |
| Raw milk | Total: | 2026967 | 2037414 | 10212.87 | 6272.15 | 121.53 | 166.82 | 1660 | 2464 |
|  | Mean: | 67565.57 | 67913.80 | 340.43 | 209.07 | 4.05 | 5.56 | 55.33 | 82.13 |
|  | SD: | 4858.92 | 5149.81 | 66.05 | 109.90 | 1.04 | 1.12 | 9.06 | 39.65 |
| Sayram Ketteki | Total: | 2065609 | 2019406 | 8085.1 | 4199.93 | 75.61 | 101.91 | 561 | 1598 |
|  | Mean: | 68853.63 | 67313.53 | 269.50 | 140.00 | 2.52 | 3.40 | 18.70 | 53.27 |
|  | SD: | 3845.32 | 4192.43 | 38.33 | 94.31 | 0.60 | 2.05 | 8.85 | 32.75 |

The rarefaction curves, which can predict the total number of species and relative abundance of each species in a sample at a given sequencing depth indicated that when the prokaryotic sequencing volume was 40,000, the bacterial sequence of each sample had not entered the plateau period (Fig 1A), and Shannon index curve (Fig 1B) indicated that all samples were still sequencing. When the amount reached 10,000, all OTUs were saturated. Although continued sequencing might discover new phylotypes, at this level, the bacterial diversity in the sample had already been captured. All the eukaryotic sequences at 10,000 had entered a gentle state, indicating that the sequencing volume of this experiment can truly reflect the diversity of fungi in the samples (Fig 1C).

**Fig 1.**
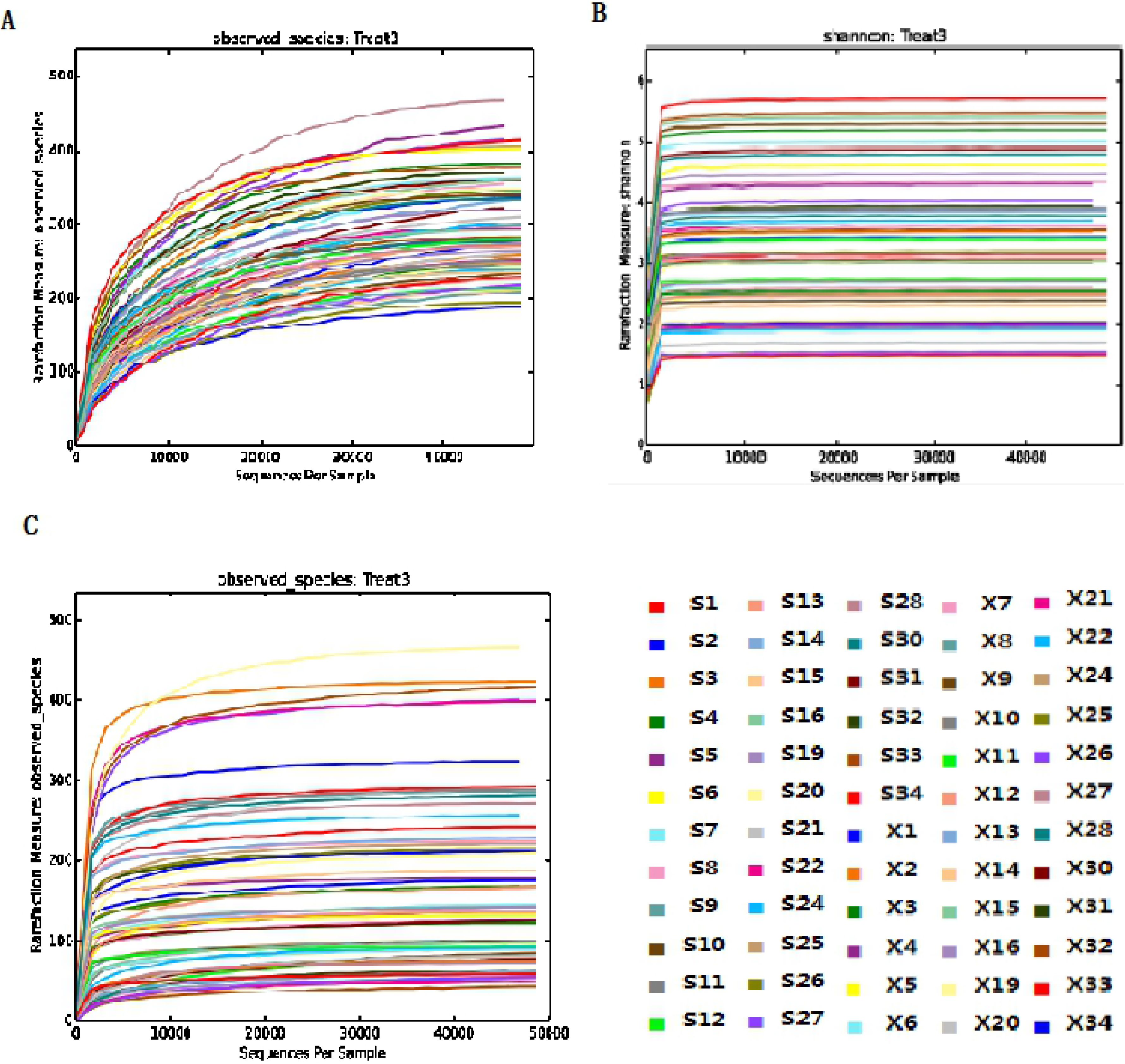
Bacterial rarefaction (A), Shannon diversity (B) and fungal rarefaction (C) curves of populations of sixty samples. Different colored lines represent different samples.

### 3.2 Comparison of bacterial and fungal community structure between RM and SK

The contribution rates of the first principal component of the sample bacteria and fungi were 61.86% and 37.29%, respectively, and the contribution rates of the second principal component were 10.4% and 21.3% (Fig 2). The closer the distance between the samples in the PCA diagram indicated the closer the composition of the two microbial communities. The SK and RM samples were separated on both sides, demonstrating that there was a significant difference in the bacterial group structure between the SK and the RM. The SK samples were almost stacked in a straight line, indicating that these samples had a high similar bacterial flora (Fig 2A). Simultaneously, the scattered distribution of bacterial communities in RM samples revealed that there were large differences between samples. The fungal samples (Fig 2B) showed a large aggregation and small dispersion. Most RM and SK samples were highly aggregated and overlapped, suggesting that these samples might have the similar fungal community, and a small part of the SK samples be scattered. This indicated that there was a difference in the fungal community among these samples.

**Fig 2.**
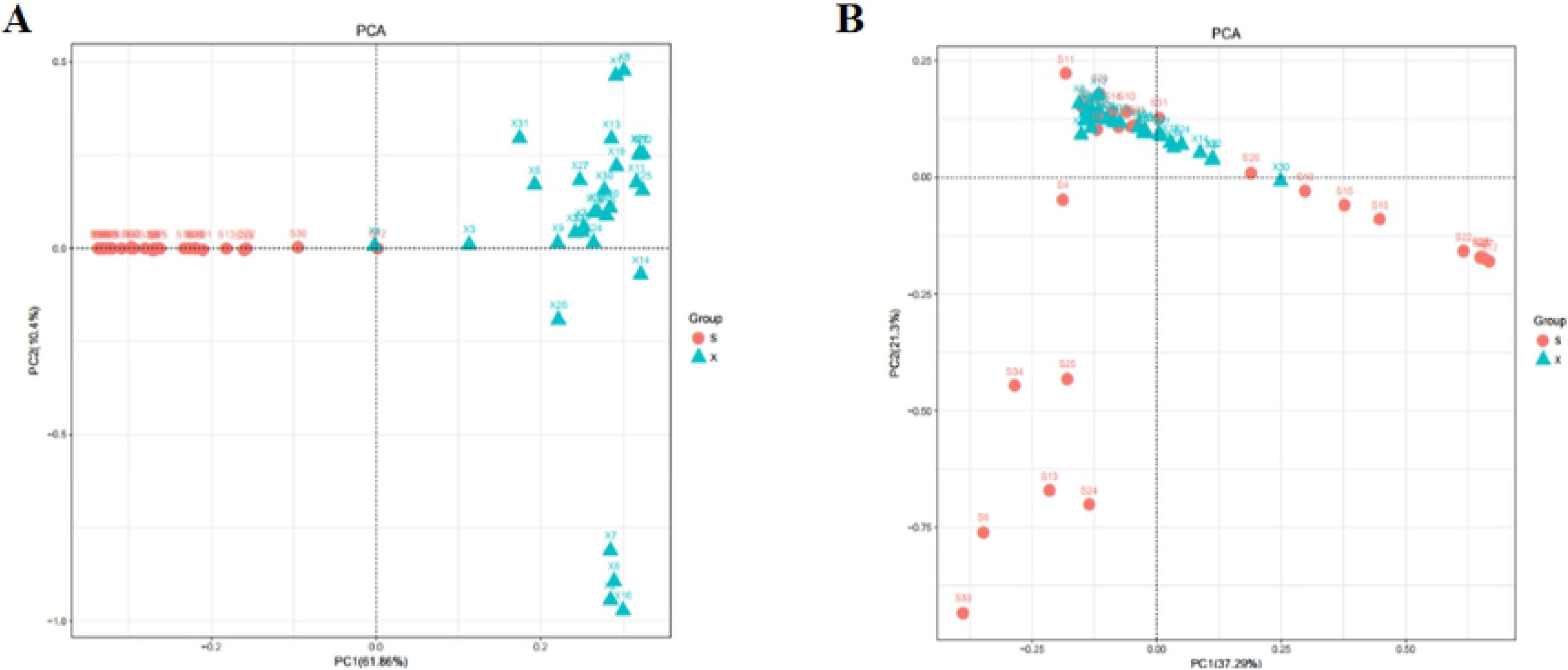
Principal component analysis (PCA) of bacterial (A) and fungal (B) community. Red circle and green triangle represent samples of Sayram Ketteki and raw milk, respectively.

### 3.3 Bacterial and fungal composition of RM and SK

A total of 15 bacteria phyla were detected in 60 samples. Firmicutes (68.50%), Proteobacteria (29.50%), and Deinococcus-Thermus (1.10%) were the dominant phyla, accounting for 99.1% of the total OTUs, while the Bacteroidetes, Actinobacteria, Cyanobacteria, Planctomycetes, Fusobacteria, Acidobacteria, Patescibacteria, Armatimonadetes, Verrucomicrobia, Elusimicrobia and Epsilonbacteraeota were observed simultaneously, indicating that the contents of them were very low, only accounting for 0.001%-0.006% of the total (Fig 3A).

**Fig 3.**
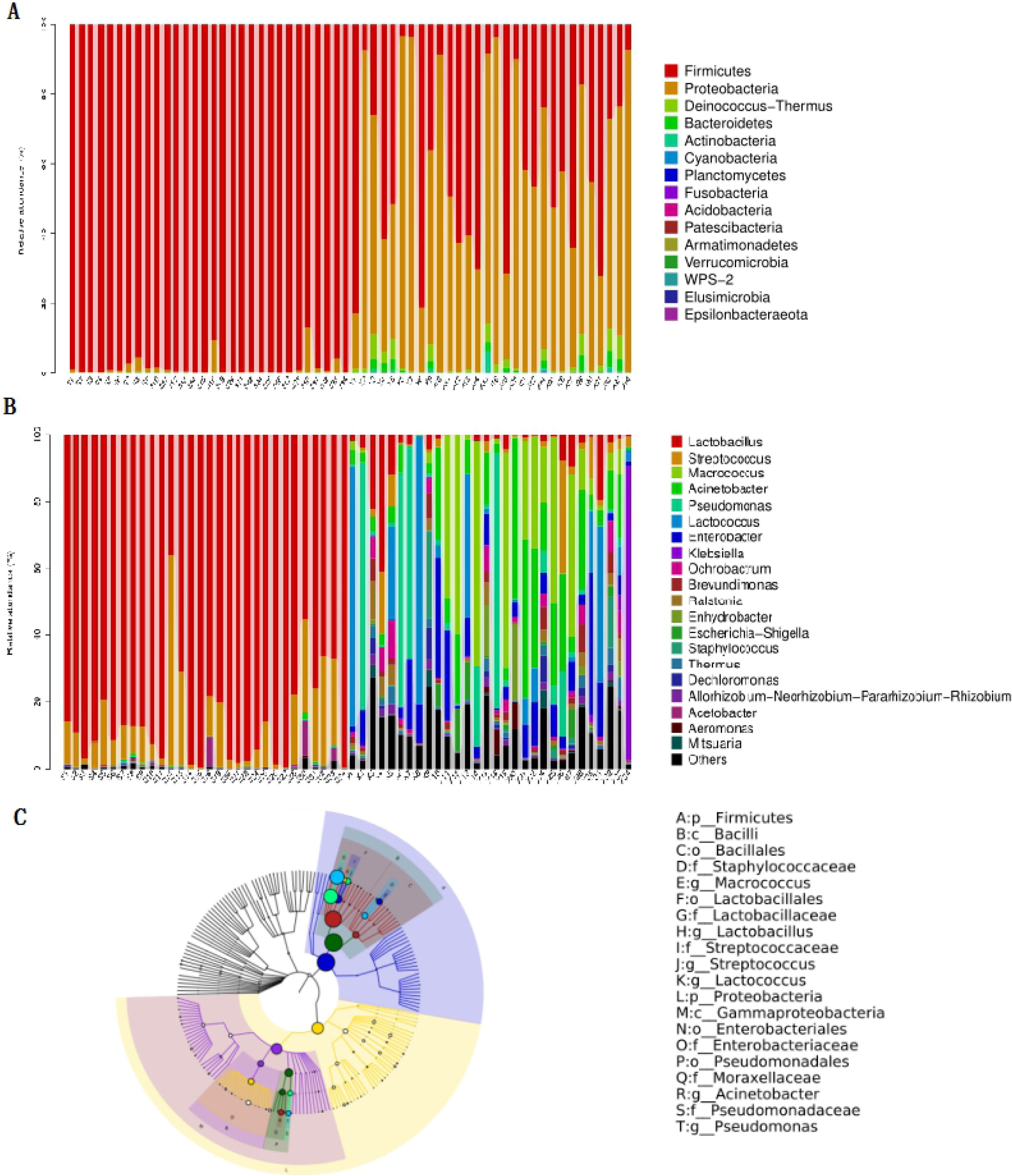
Relative abundances of bacteria at phylum (A) and genera (B) levels in each samples and GraPhlAn hierarchical tree map (C).

A total of 218 genera of bacteria were detected, and there were 11 genera with abundance greater than 1%, including *Lactobacillus* (45.30%), *Streptococcus* (8.00%), *Macrococcus* (7.70%), *Acinetobacter* (6.70%), *Pseudomonas* (6.00%), *Lactococcus* (5.60%), *Enterobacter* (4.50%), *Klebsiella* (1.60%), *Ochrobactrum* (1.40%), *Brevundimona* (1.10%) and *Ralstonia* (1.00%) (Fig 5B). Moreover, it revealed that the *Macrococcus*, *Acinetobacte*r, *Pseudomonas* and *Lactococcus* were the dominant genera in RM, with proportion being 15.3%, 13.1%, 11.9% and 11.15%, respectively. *Lactobacillus* and *Streptococcu*s in SK were the dominant bacterial genera, accounting for 98.1% (Table 2 and Fig 3C). The content of *Lactobacillus* in these samples ranged from 36.4% to 99.7%, which might be closely related to the sampling location, processing environment and other factors. The OTUs content of *Lactobacillus* and *Streptococcus* increased significantly after processing, while other 18 bacterial species decreased dramatically. For example, the OTUs content of *Staphylococcus* dropped from 1.66% to 0, presuming that the *staphylococcus* was completely killed after heating treatment.

**Table 2.** Analysis of OTU Percentage of the top 20 genera

| Genera of Bacteria | OTU Percentage (%) |  | Genera of Fungi | OTU Percentage (%) |  |
| --- | --- | --- | --- | --- | --- |
|  | RM | SK |  | RM | SK |
| *Lactobacillus* | 5.1067±8.727 | 85.5967±14.883 | *Candida* | 13.400±11.517 | 28.9233±35.865 |
| *Streptococcus* | 3.5533±7.047 | 12.4667±13.646 | *Kluyveromyces* | 0.4833±1.007 | 14.8000±29.498 |
| *Macrococcus* | 15.3033±22.684 | 0.0033±0.018 | *Aspergillus* | 6.2967±4.311 | 3.9433±11.780 |
| *Acinetobacter* | 13.1400±13.859 | 0.2033±0.365 | *Lecanicillium* | 4.9467±4.301 | 2.9133±4.909 |
| *Pseudomonas* | 11.9867±24.274 | 0.0633±0.119 | *Mucor* | 2.1267±5.799 | 3.6867±16.573 |
| *Lactococcus* | 11.1467±22.138 | 0.0200±0.109 | *Penicillium* | 2.9167±4.584 | 0.8433±1.692 |
| *Enterobacter* | 8.9133±10.551 | 0.0567±0.122 | *Clavispora* | 1.7633±8.781 | 1.9900±10.336 |
| *Klebsiella* | 3.2467±16.039 | 0.0067±0.0365 | *Malassezia* | 2.6133±2.385 | 1.0200±1.941 |
| *Ochrobactrum* | 2.7500±3.114 | 0.0533±0.073 | *Yarrowia* | 0.6500±2.244 | 2.9000±15.188 |
| *Brevundimonas* | 2.0067±2.398 | 0.0933±0.151 | *Cladosporium* | 2.7967±3.484 | 0.6533±1.147 |
| *Ralstonia* | 2.0433±2.861 | 0.0367±0.096 | *Thermoascus* | 2.9467±3.831 | 0.4533±0.807 |
| *Enhydrobacter* | 1.7067±4.902 | 0.0067±0.025 | *Simplicillium* | 1.8167±1.786 | 1.1433±1.679 |
| *Escherichia* | 1.6700±3.396 | 0.0067±0.025 | *Verticillium* | 1.7100±1.453 | 0.9067±1.689 |
| *Staphylococcus* | 1.6633±5.496 | 0.0000±0.000 | *Suillus* | 2.3433±4.163 | 0.0267±0.146 |
| *Thermus* | 1.4267±1.737 | 0.0367±0.056 | *Pichia* | 0.8700±0.763 | 1.4600±2.719 |
| *Dechloromonas* | 1.2000±2.364 | 0.0033±0.018 | *Cutaneotrichosporon* | 1.5433±3.325 | 0.7800±1.825 |
| *Allorhizobium* | 0.9933±1.145 | 0.0100±0.031 | *Trichosporon* | 1.8533±9.383 | 0.2067±0.549 |
| *Acetobacter* | 0.0167±0.046 | 0.0073±0.024 | *Rhodotorula* | 1.8100±8.485 | 0.1500±0.395 |
| *Aeromonas* | 0.7133±1.933 | 0.0002±0.001 | *Thermomyces* | 1.8067±2.833 | 0.0467±0.104 |
| *Mitsuaria* | 0.6733±1.441 | 0.0001±0.001 | *Wickerhamomyces* | 1.4667±2.552 | 0.0700±0.195 |

A total of 12 fungal phyla were detected successfully (Fig 4A). Ascomycota (79.90%), Basidiomycota (10.20%) and Mucoromycota(3.40%), with an average abundance value greater than 1% were absolutely dominant, accounting for 93.5% of the total, while the Mortierellomycota, Rozellomycota, Olpidiomycota, Entomophthoromycota, Chytridiomycota, Glomeromycota, Zoopagomycota, GS19 and other unidentified phylum, had very small content, only accounting for 0.1%-0.5% of the total OTUs. There were 495 fungal genera detected, and there were 18 genera with an average abundance value greater than 1%, including *Candida* (21.20%), *Kluyveromyces* (7.60%), *Aspergillus* (5.10%), *Lecanicillium* (3.90%), *Muco*r (2.90%), *Penicillium* (1.90%) *Clavispora* (1.90%) *Malassezia* (1.80%), *Yarrowia* (1.80%), *Cladosporium* (1.70%), *Thermoascus* (1.70%), *Simplicillium* (1.50%), *Verticillium* (1.30%), *Suillus* (1.20%), *Pichia* (1.20%), *Cutaneotrichosporon* (1.20%), *Trichosporon* (1.00%) and *Rhodotorula* (1.00%) (Fig 4B). It indicated that *Candida* had the highest content in the two types of samples (Table 2 and Fig 4C). After fermentation, the genera contents of *Candida* and *Kluyveromyces* increased significantly. The OTUS contents of *Mucor*, *Clavispora*, *Yarrowia* and *Pichia* increased slightly, and other 14 fungal genera decreased significantly.

**Fig 4.**
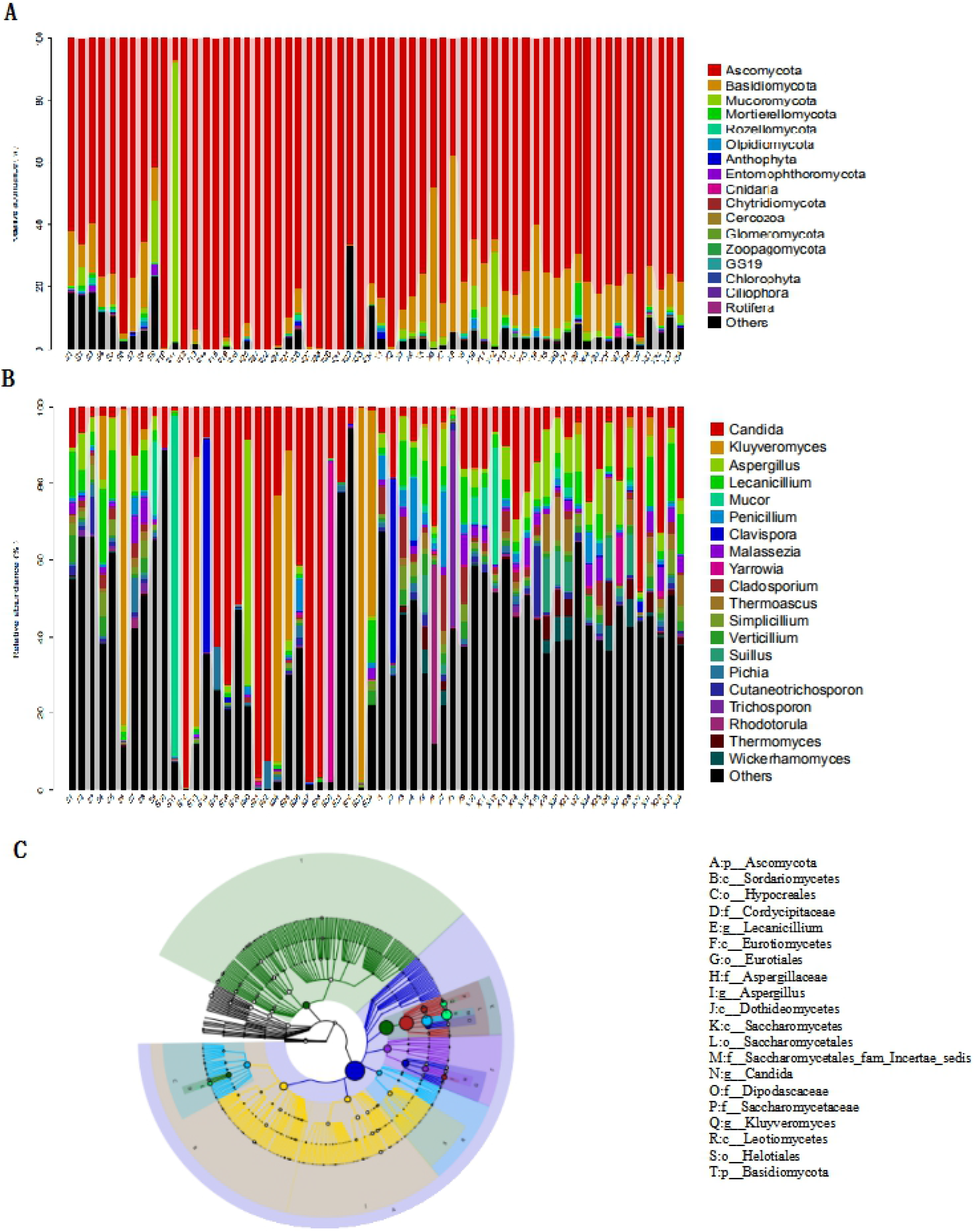
Relative abundances of fungi at phylum (A) and genera (B) levels in each samples and GraPhlAn hierarchical tree map (C).

In all, we found that the OTUs of predominant bacterial genera in SK was higher and obvious, and the OTUs of fungal genera were relatively uniform and small in both two types of dairy products (Table 2).

### 3.4 Spearman association network analysis of dominant genera interaction

The association network of dominant bacteria was constructed by using Cytoscape, and the most of the bacterial relationships were positive (Fig 5). The *Lactococcus* was positively correlated with more than 20 species of bacteria for example *Klebsiella*, *Staphylococcus*, *Enterobacter* and so on, while the *Lactobacillus* was negatively correlated with more than 20 species of bacteria such as *Enterococcus*, *Lactococcus* and *Streptococcus.* It suggested that the lactic acid bacteria might ferment lactose to produce lactic acid, which created an acidic environment. The acid tolerance of *Lactobacillus* was higher than other bacteria, especially *Lactococcus.* This also explained the high abundance of *Lactobacillus* in yogurt, and higher abundance of *Lactococcus* in RM.

**Fig 5.**
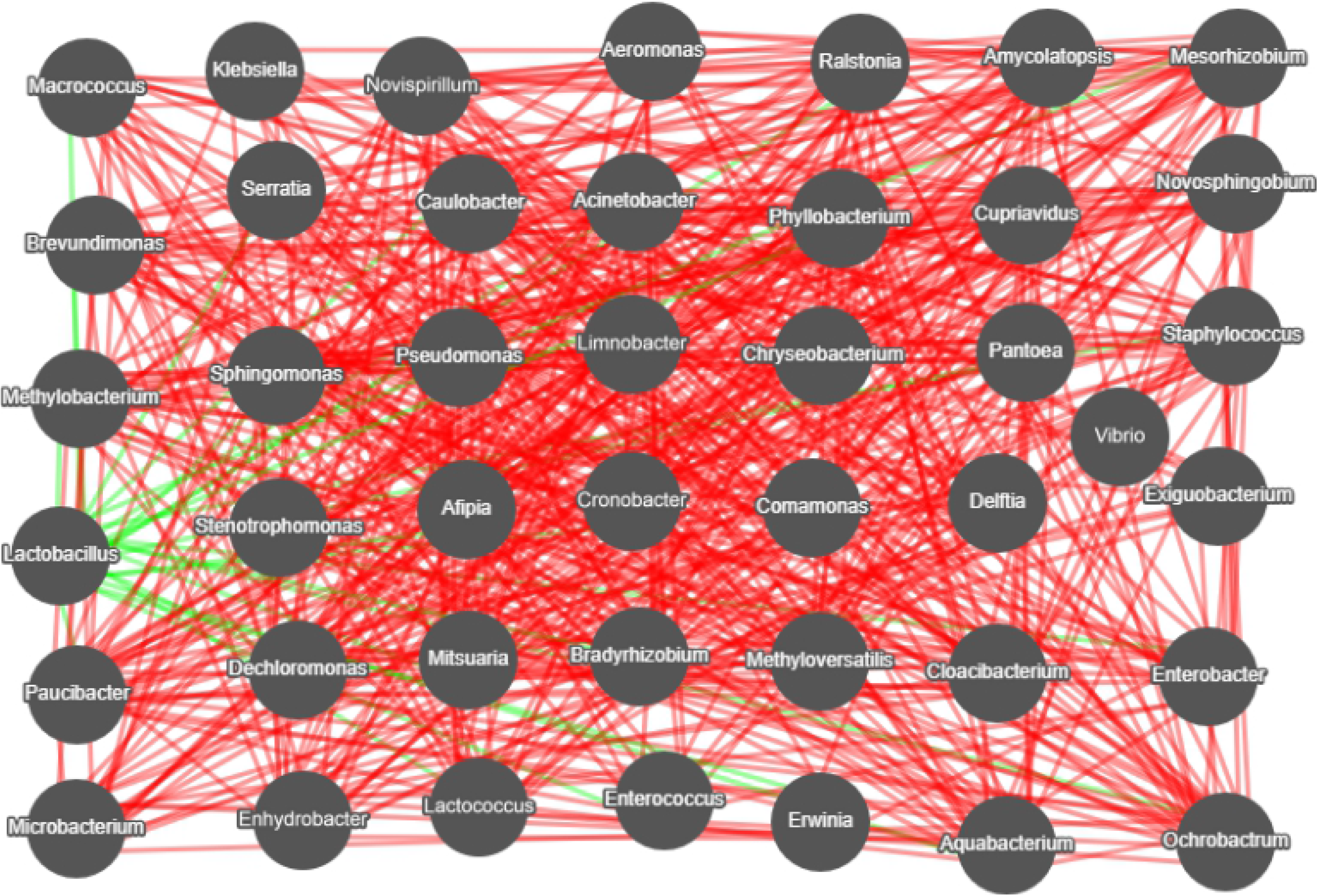
Dominant of top 50 bacterial genera association network diagram. The red and green lines indicates the positive and negative relationships, respectively.

### 3.5 Prediction of flora metabolism function

Using the PICRUSt, the prediction of bacterial metabolic functions can be achieved by comparing the existing 16SrRNA gene sequencing data with the microbial reference genome database with known metabolic functions. The composition of the KEGG two-level functional prediction information obtained from RM and SK was basically similar, but the abundance was obviously different (Fig 6A).

Bacterial community with higher abundance of RM was involved in the two classifications of carbohydrate metabolism and amino acid metabolism. SK had the largest variety of flora for carbohydrate metabolism. Venn diagram of common functional groups showed that although there were obvious differences in the composition of the microbial community, the functional groups in each type of sample were basically similar (Fig 6B).

**Fig 6**. PICRUSt predicted KEGG second level distribution map (A) and Venn diagram of common functional groups (B).

## 4. Discussion

The multiple microorganism and species diversity exist in RM[38] and traditional fermented dairy products[39, 40], and they play a crucial role in the formation of flavor. Studies have shown that the main source of the flavor substance acetaldehyde in yogurt is the synthesis of threonine by threonine aldolase secreted by the *Bulgaria* subspecies, and *Streptococcus thermophilus* can metabolize citric acid to produce diacetyl during yogurt fermentation and storage [41, 42]. There are various and popular fermentation dairy products in Xinjiang rural area, such as Kurut, Shubat, hence, there is no doubt that their particular flavor has a lot to do with the microbial polymorphism. However, the microbial diversity of these dairy products remain elusive. SK is one of the beloved yogurt and its microbial diversity has been not investigated yet, so, it’s the first time to study its microbial polymorphism in the current study.

Previous studies revealed that *Lactobacillus* is the main flora in cow’s milk [43, 44]. In the current study, the *Macrococcus*, *Acinetobacter*, *Pseudomonas* and *Lactococcus* were the dominant bacterial genera in RM, accounting for 51.5% of the total sequences, but the proportion of each sample is different. The *Lactobacillus* and *Streptococcus* were the dominant bacterial genera in SK, which are the similar to other studies [45]. At the fungal level, *Candida* was the absolutely advantageous genus in both two types of dairy products, while *Aspergillus* and *Kluyveromyces* were the subsequent dominant genera in RM and SK samples, which differed with others study [46]. The *Kluyveromyces marxianus* used in fermented food had the characteristics of weak alcohol and good aroma production, and could increase the content of free fatty acids and flavor substances in yogurt [47, 48]. Maybe, that’s why the flavor of SK is different from the other yogurts.

There were huge differences in the types and contents of microbial flora between samples. The most SK samples contained more than 50% *Lactobacillus* and about 10% *Streptococcus*, but in S12, the contents of *Lactobacillu*s and *Streptococcus* were 36.4% and 62.9%, respectively. At fugal level, *Candida* was the dominant genus which the average content in RM was 13.4%. In sample X12, *Candida* accounted for only 0.5%, while *Trichosporon* accounted for 51.5%. That may be caused by sampling. In the future, samples should be collected on a larger scale for further research.

In all, our findings partially explore the microbial diversity of RM and SK in south Xinjiang and contribute to providing a basis for industrialization SK. Besides, the microbial functional prediction can be helpful to search for new probiotics.

## Acknowledgments

We would like to thank Dr Leilei Zhan for their helpful discussions and critical reading of the manuscript.

